# Systemic tumor-cell reprogramming converts solid tumors into *in situ* biofactories for immunotherapy

**DOI:** 10.64898/2026.09.26.754636

**Authors:** Shihui Wang, Qing Xu, Mengyao Wu, Jiarong Duan, Qin Gao, Haimin Ma, Wei Li, Tianqing Zheng

## Abstract

A central challenge in oncology is delivering potent drugs to tumors without harming healthy tissues. We therefore asked whether tumor cells themselves could be genetically programmed to produce therapeutic molecules directly within the tumor. We present LENTRA (LENtiviral Tumor-Reprogramming Activator), an antigen-targeted lentiviral platform that enables selective transduction and genetic modification of tumor cells *in vivo*. Following intravenous administration, LENTRA achieves tumor-restricted expression of immune checkpoint inhibitors, cytokines, or therapeutic enzymes in multiple murine solid tumor models. Local production of anti-CTLA-4 or IL-12 triggered potent antitumor responses while avoiding systemic toxicities. Tumor-restricted secretion of sialidase enhanced dendritic cell and T cell infiltration, and reprogrammed tumor-associated macrophages toward an immunostimulatory phenotype, resulting in tumor regression and durable immune memory, including in tumors with heterogeneous antigen expression. In patient-derived tumor explants, LENTRA mediated efficient transduction, immune remodeling, and reduced tumor viability. By engineering tumors to produce therapeutic proteins, LENTRA converts resistant tumors into autonomous therapeutic sources and active immune hubs, achieving local TME reprogramming and systemic antitumor immunity with an improved safety profile.

## Introduction

Many immunomodulatory agents can induce strong antitumor responses, yet their clinical application is constrained by systemic toxicity or by poor drug penetration into the tumor microenvironment (TME)(1–4). One solution is local administration, which can achieve high intratumoral drug concentrations; yet it is practical only for accessible lesions and does not address metastatic disease(5, 6). An alternative strategy is systemic gene delivery, enabling the tumor itself to produce therapeutic molecules locally(7). Such disease-site programming could offer the advantages of systemic delivery while achieving sustained, spatially confined therapeutic activity at the tumor site.

Recent work has shown that cells can be reprogrammed *in vivo* using nucleic acids or viral vectors. LNP-mRNA platforms, for example, can transiently turn tumors into protein-producing sites, but most such applications require intratumoral injection; systemically infused LNPs tend to accumulate in the liver(8–11). Oncolytic viruses offer another route for intratumoral protein production, yet their activity is tied to replication and lysis, and systemic administration has been hampered by host immunity and hepatic sequestration(12, 13).

Receptor-retargeted lentiviral vectors have been extensively developed for cell-selective gene delivery, with recent efforts focused on *in vivo* CAR-T generation(14–20). Tumor-directed lentiviral targeting has also been explored. Previous studies demonstrated selective transduction of tumor cells *in vitro* and, in some cases, tumor-selective transgene expression following systemic administration, although biodistribution varied substantially across vector designs(15–17). Intravenously administered cetuximab-targeted lentiviral vectors, for example, accumulated predominantly in the liver and spleen, with comparatively lower accumulation in tumors(16).

These studies showed that lentiviral vectors can be redirected toward tumor-associated receptors, but left open whether systemic tumor-directed delivery could be developed into a modular strategy for programming tumor cells as sustained local producers of therapy. Such an approach would require sufficient selectivity to confine potent payload production to the tumor while retaining the versatility to support distinct therapeutic mechanisms.

Here, we address this question with LENTRA (LENtiviral Tumor-Reprogramming Activator), a tumor-retargeted lentiviral platform designed to convert solid-tumor cells into *in situ* therapeutic biofactories (**Fig. 1a**). Tumor-selective transduction is achieved by pseudotyping lentiviral envelopes with tumor-associated antigen (TAA)-binding proteins and a tropism-deficient, fusion-competent VSV-G mutant(21–24). LENTRA drives tumor-restricted expression of checkpoint inhibitors, cytokines, or enzymatic effectors, concentrating therapeutic activity within the TME while limiting systemic exposure. In murine models and patient-derived tumor explants, this localized production remodels the tumor ecosystem and elicits potent antitumor activity—including tumor regression *in vivo* and direct killing *ex vivo*. These findings establish systemic tumor-cell programming as a modular therapeutic strategy that converts the disease tissue from a passive treatment target into a local source of therapy.

**Fig. 1.**
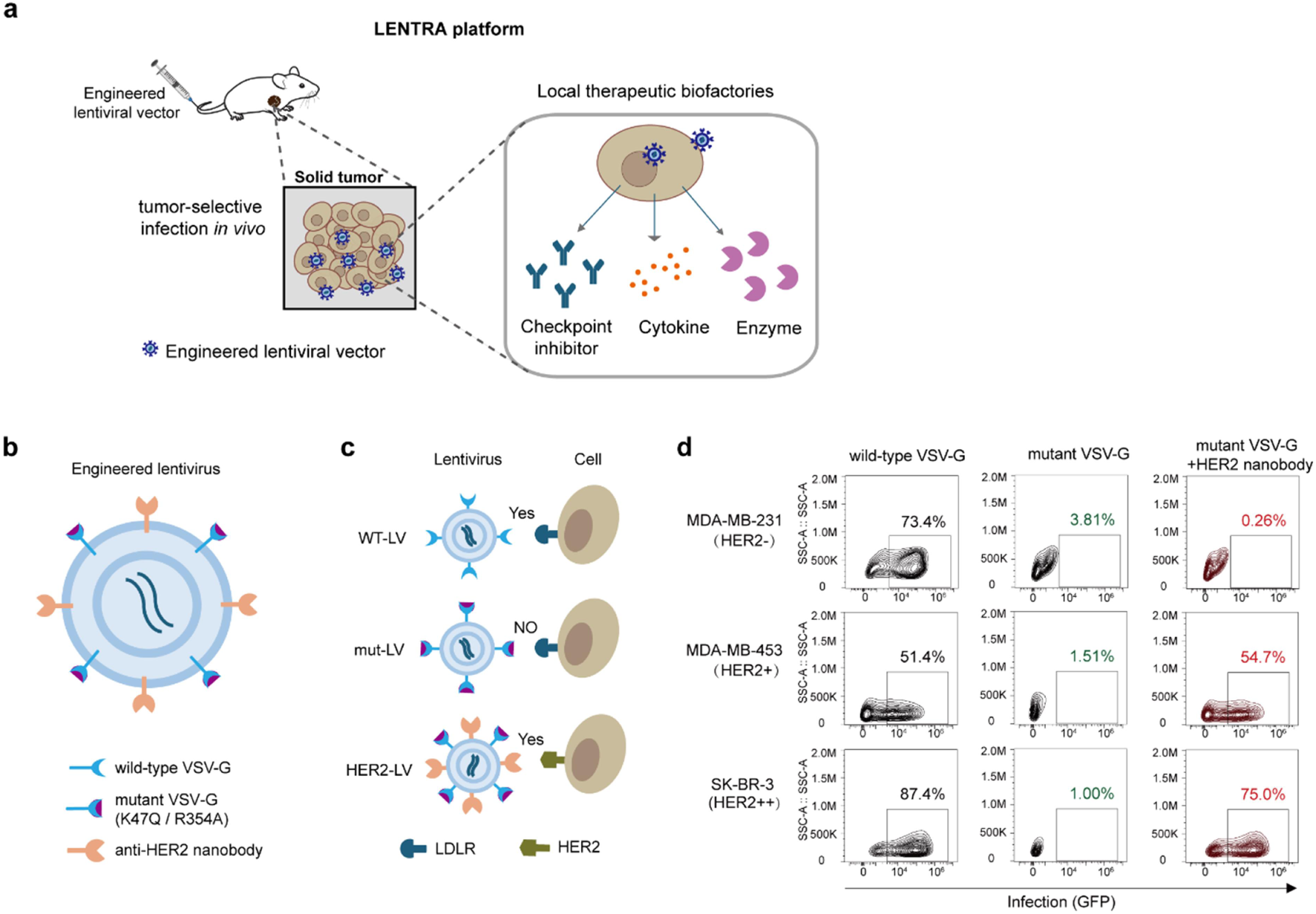
Design of the LENTRA platform. (a) Schematic of the LENTRA platform developed in this study, which enables genetic engineering of tumors *in vivo* following intravenous administration. (b) Design of the HER2-targeted lentiviral vector. Lentiviral particles were pseudotyped with a VSV-G mutant (K47Q/R354A) and a membrane-anchored anti-HER2 nanobody. (c) The three pseudotyped vectors used in this study. WT-LV carries wild-type VSV-G; mut-LV carries mutant VSV-G (K47Q/R354A) alone; HER2-LV carries both mutant VSV-G (K47Q/R354A) and the membrane-anchored anti-HER2 nanobody. (d) HER2-negative (MDA-MB-231) and HER2-positive (MDA-MB-453, SK-BR-3) cells were infected with GFP-encoding WT-LV, mut-LV, or HER2-LV. GFP expression was analyzed by flow cytometry 48 h post-infection.

## Results

### Engineering a tumor-targeted lentiviral platform

To create lentiviral vectors that recognize tumors rather than the ubiquitous LDL receptor, we engineered a platform that decouples target recognition from membrane fusion. Conventional lentiviral vectors are pseudotyped with VSV-G, a glycoprotein that does two things: it binds LDLR to initiate endocytosis, and it fuses membranes within endosomes to release the viral genome(23). Because LDLR is ubiquitously expressed on mammalian cells, VSV-G– pseudotyped lentiviral vectors lack cell selectivity(23). To redirect tropism to tumors, we combined two envelope components: a VSV-G mutant (K47Q/R354A)(21–24) that retains fusion capacity but cannot bind LDLR, and a membrane-anchored anti-HER2 nanobody(25) that recognizes HER2, a tumor-associated antigen overexpressed on many epithelial malignancies (**Fig. 1b** and **fig. S1**).

To test this design, we produced GFP-encoding lentiviruses with three distinct pseudotypes: wild-type VSV-G (WT-LV, canonical broad tropism)(23), mutant VSV-G alone (mut-LV, loss-of-infection control)(23), and a combination of mutant VSV-G with the anti-HER2 nanobody (HER2-LV) (**Fig. 1c** and **fig. S1**). Infection of a panel of breast cancer cell lines that differ in HER2 expression showed that WT-LV transduced both HER2-negative (MDA-MB-231) and HER2-positive (MDA-MB-453, SK-BR-3) cells with similar efficiency. mut-LV showed near-complete loss of infectivity, confirming that native tropism had been abolished. HER2-LV, however, selectively infected HER2-positive cells (>50% GFP⁺ in MDA-MB-453 and SK-BR-3) while producing only background signal in HER2-negative cells (**Fig. 1d**). Thus, HER2-LV redirected lentiviral tropism to HER2-positive cells.

To verify that HER2-LV infection requires HER2 engagement, we performed a competitive blockade assay. HER2-positive MDA-MB-453 cells were pre-incubated with increasing concentrations (0–163 nM) of a HER2-neutralizing antibody before exposure to mCherry-encoding HER2-LV. Transduction efficiency decreased in a dose-dependent manner, indicating that the antibody blocks viral entry by competing with HER2-LV for HER2 binding. (**Fig. 2a**).

**Fig. 2.**
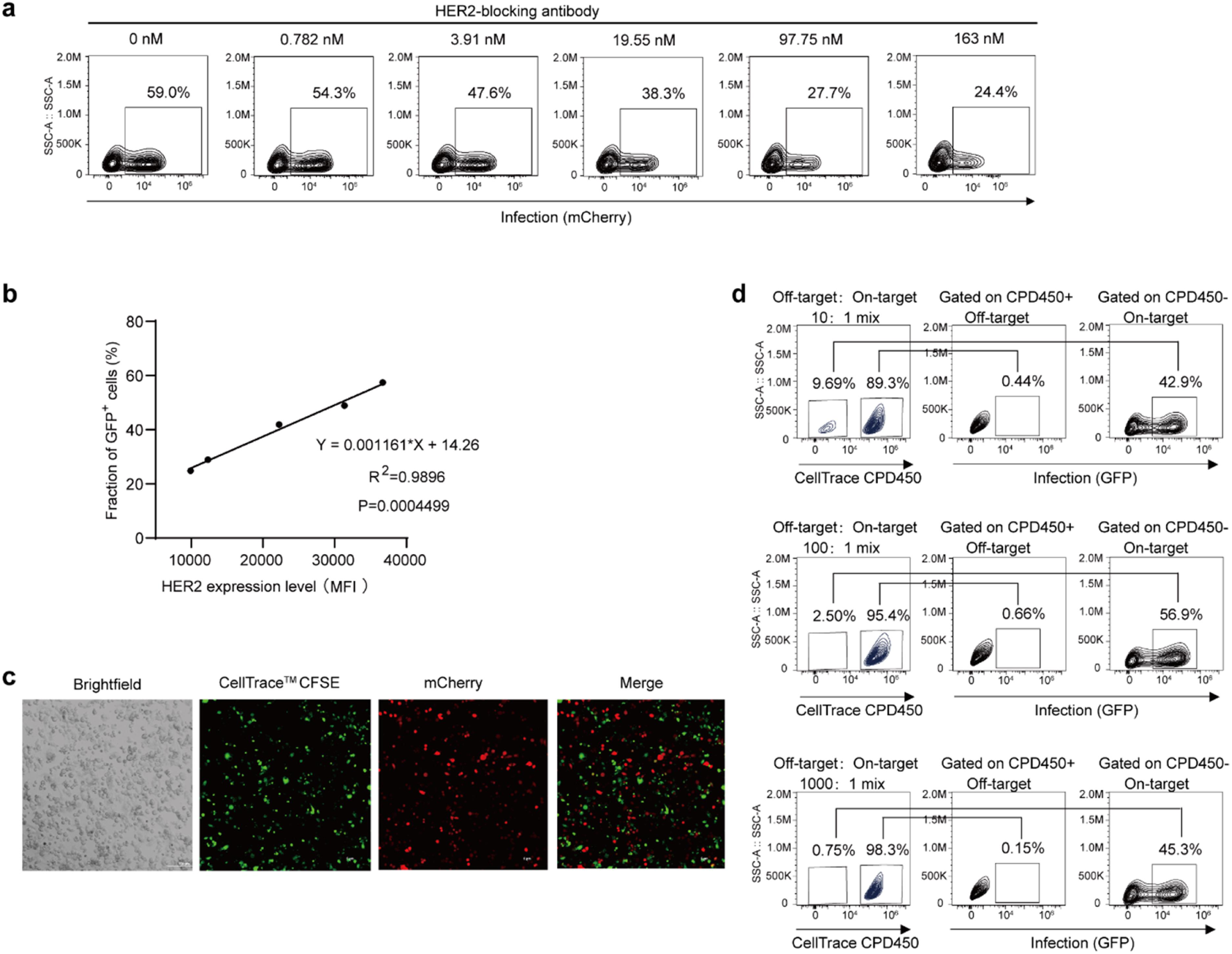
LENTRA transduces HER2-positive cells with high specificity and sensitivity. (a) MDA-MB-453 cells were pre-incubated with increasing concentrations of a HER2-blocking antibody before exposure to mCherry-encoding HER2-LV. mCherry expression was analyzed by flow cytometry 48 h post-infection. (b) HEK293T cells expressing graded HER2 levels were infected with GFP-encoding HER2-LV and analyzed by flow cytometry 48 hours later. The fraction of GFP⁺ cells showed a linear correlation with HER2 expression level. (c) CFSE-labeled HER2-negative MDA-MB-231 cells were co-cultured with unlabeled HER2-positive MDA-MB-453 cells, infected with mCherry-encoding HER2-LV, and imaged by confocal microscopy 48 h post-infection (scale bar, 0.2 mm). (d) CPD450-labeled HER2-negative MDA-MB-231 cells were mixed with HER2-positive MDA-MB-453 cells at ratios of 10:1, 100:1, or 1000:1, infected with GFP-encoding HER2-LV, and analyzed by flow cytometry 48 h post-infection.

We next examined whether transduction efficiency scales with HER2 surface density. HEK293T cells were engineered to express graded HER2 levels, then infected with GFP-encoding HER2-LV. Transduction efficiency correlated strongly with HER2 mean fluorescence intensity (MFI) (R² > 0.98), indicating that transduction output scales quantitatively with antigen density (**Fig. 2b**).

To evaluate selectivity in mixed populations, we co-cultured CFSE-labeled HER2-negative MDA-MB-231 cells with unlabeled HER2-positive MDA-MB-453 cells at a 1:1 ratio and added mCherry-encoding HER2-LV. Confocal microscopy performed 48 hours after infection showed mCherry expression exclusively in HER2-positive cells, with no detectable signal in CFSE-labeled HER2-negative cells (**Fig. 2c**). We next tested the system under extreme target dilution, mixing HER2-positive cells with HER2-negative cells at ratios of 1:10, 1:100, and 1:1000. After infection with GFP-encoding HER2-LV, flow cytometry showed that even at 0.1% target frequency, HER2-LV maintained high selectivity (**Fig. 2d**). This sensitivity is important for systemic delivery, where target cells are vastly outnumbered by normal tissues.

### LENTRA enables selective transduction of tumors *in vivo*

Having confirmed stringent selectivity *in vitro*, we tested whether HER2-LV could achieve tumor-specific transduction after intravenous administration (**Fig. 3a**). BALB/c mice bearing orthotopic EMT6-HER2 tumors in the mammary fat pad received a single injection of GFP-encoding HER2-LV when tumors reached ∼100 mm³. Ten days later, ex vivo imaging of harvested organs showed strong GFP signal confined to the tumors, with no detectable signal above background in heart, liver, spleen, lung, or kidney (**Fig. 3b**).

**Fig. 3.**
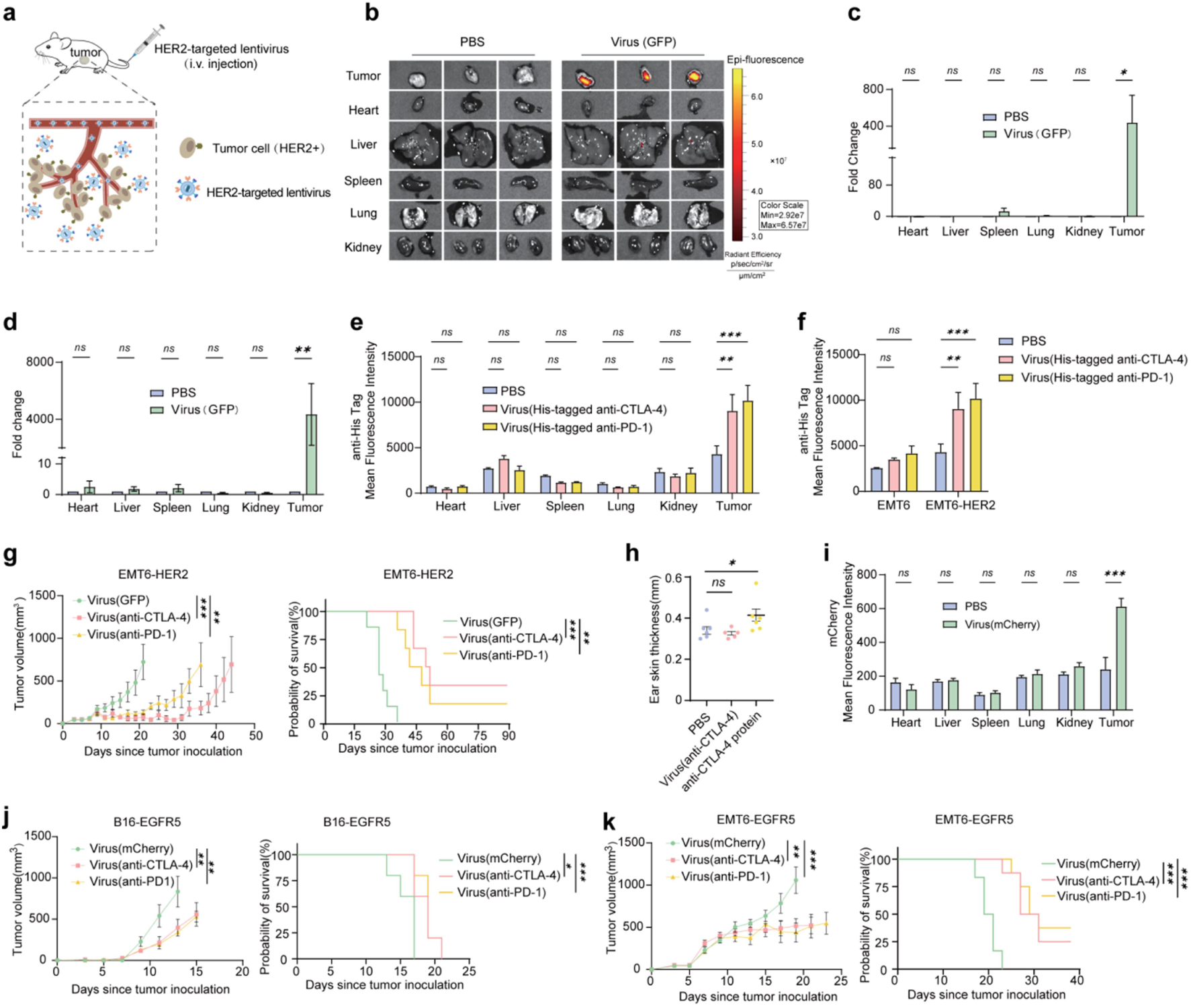
LENTRA converts tumors into local factories for checkpoint blockade. (a) Schematic of HER2-LV-mediated transgene delivery to HER2-positive tumors. (b) BALB/c mice bearing orthotopic EMT6-HER2 tumors received an intravenous injection of GFP-encoding HER2-LV or PBS (n = 3 per group). Tumors and organs were harvested 10 days later and imaged on an IVIS Spectrum system. (c) GFP copy number in genomic DNA from tumors and normal tissues was measured by qPCR. (d) GFP mRNA levels in tumors and normal tissues were measured by RT-qPCR and normalized to housekeeping genes. (e) Mice bearing EMT6-HER2 tumors were treated intravenously with PBS or HER2-LV encoding His-tagged anti-CTLA-4 or anti-PD-1. After ten days, single-cell suspensions prepared from tumors and normal tissues were stained with Alexa Fluor™ 647 anti-His tag antibody, and analyzed by flow cytometry. (f) His-tag expression in tumors isolated from EMT6 or EMT6-HER2 tumor-bearing mice treated as in (e). (g) Tumor growth and survival of EMT6-HER2-bearing mice treated with GFP-encoding HER2-LV (n=7), anti-CTLA-4-encoding HER2-LV (n=5), or anti-PD-1-encoding HER2-LV (n=6). (h) DNFB-induced contact hypersensitivity mice bearing EMT6-HER2 tumors were injected intravenously with anti-CTLA-4 protein or anti-CTLA-4-encoding HER2-LV. Ear thickness was measured after DNFB challenge. (i) Mice bearing B16-EGFR5 tumors received PBS or mCherry-encoding EGFR-LV. Single-cell suspensions prepared from tumors and normal tissues were analyzed for mCherry expression by flow cytometry six days later. (j) Tumor growth and survival of mice bearing B16-EGFR5 tumors treated with mCherry-, anti-CTLA-4-, or anti-PD-1-encoding EGFR-LV (n=5). (k) Tumor growth and survival of mice bearing EMT6-EGFR5 tumors (n=6-8) treated with mCherry-, anti-CTLA-4-, or anti-PD-1-encoding EGFR-LV. ***p < 0.0001; **p < 0.01; *p < 0.05 by Student’s t-test for (c, d, e, f, i); by log-rank (Mantel-Cox) test for survival curves in (g, j, k); by one-way ANOVA with Tukey’s post-hoc test for (h); and by two-way ANOVA with Tukey’s post-hoc test for tumor growth curves in (g, j, k).

We further assessed the *in vivo* selectivity of HER2-LV by analyzing viral DNA integration and transgene expression. Quantitative PCR (qPCR) on genomic DNA confirmed that integrated GFP copies were enriched in tumors relative to normal tissues (**Fig. 3c**). To assess functional transgene expression, we performed reverse-transcription qPCR (RT-qPCR) for GFP mRNA. Tumors exhibited robust GFP transcript levels, while normal tissues showed no significant difference from PBS-injected controls (**Fig. 3d**). Thus, HER2-LV achieves tumor-restricted transduction and transgene expression after intravenous administration.

### LENTRA converts tumors into local factories for checkpoint blockade

Having shown tumor-restricted delivery, we asked whether HER2-LV could convert tumors into local factories for therapeutic antibody production. We first tested engineering tumors to produce checkpoint inhibitors *in vivo*. BALB/c mice with established EMT6-HER2 tumors received an intravenous dose of HER2-LV encoding His-tagged anti-PD-1 or anti-CTLA-4(3, 26). Flow cytometry of tumors harvested seven days later showed that tumor cells were His-tag⁺, with no significant signal in normal tissues, confirming tumor-restricted antibody production (**Fig. 3e**). Specificity was further validated in mice bearing HER2-negative EMT6 tumors, where the same virus yielded no signal above PBS controls (**Fig. 3f**). Therapeutically, a single dose of anti-PD-1- or anti-CTLA-4-encoding HER2-LV significantly suppressed tumor growth compared with GFP-encoding virus (**Fig. 3g** and **fig. S2a**).

Beyond efficacy, an advantage of localized production is the potential to mitigate systemic immune-related adverse events (irAEs), a major limitation of checkpoint blockade(1). Systemic administration of anti-CTLA-4 antibodies can cause dose-limiting irAEs such as dermatitis, colitis, and hepatitis—reflecting broad immune activation beyond the tumor site(2). To test whether tumor-restricted anti-CTLA-4 production using HER2-LV could reduce such toxicity, we used a 2,4-dinitrofluorobenzene (DNFB)-induced contact hypersensitivity model(27). Intravenous injection of recombinant anti-CTLA-4 protein (200 µg per dose) significantly exacerbated ear swelling. By contrast, mice treated with anti-CTLA-4-encoding HER2-LV, which confines antibody production to the tumor, showed no significant exacerbation of inflammation (ear swelling comparable to the PBS group) (**Fig. 3h**), indicating a favorable safety profile.

We then tested the platform’s modularity by targeting a different tumor antigen. An EGFR-targeted lentivirus (EGFR-LV) was designed by pseudotyping with the mutant VSV-G protein (K47Q/R354A) and a membrane-anchored anti-EGFR scFv derived from cetuximab(28). In C57BL/6 mice bearing B16-EGFR5 tumors(29), which express a chimeric mouse EGFR recognized by cetuximab, intravenous injection of mCherry-encoding EGFR-LV yielded tumor-restricted transgene expression. (**Fig. 3i**). Therapeutically, EGFR-LV encoding anti-PD-1 or anti-CTLA-4 effectively suppressed tumor growth in both B16-EGFR5 and EMT6-EGFR5 mouse tumor models (**Fig. 3j,k** and **fig. S2b,c**). These results demonstrate that the LENTRA platform is generalizable across different tumor antigens and tumor types.

### Tumor-restricted IL-12 production via LENTRA bypasses systemic toxicity

Given that LENTRA enables tumor-restricted production of biologics, we reasoned that this platform might be particularly useful to deploy potent drugs that are otherwise systemically intolerable. We tested this with interleukin-12 (IL-12), a cytokine with potent antitumor activity but dose-limiting systemic toxicity that has hindered its clinical application(30, 31). Mice bearing EMT6-HER2 tumors were injected intravenously with His-tagged IL-12-encoding HER2-LV or PBS. Flow cytometry showed His-tagged IL-12 was detected in tumors, but not in healthy tissues (**Fig. 4a**). This tumor-localized IL-12 production led to complete regression of tumors in all treated mice, which remained tumor-free long-term (**Fig. 4b** and **fig. S3a**). Immunohistochemistry and flow cytometry revealed a remodeled TME, with increased T cell infiltration and reduced regulatory T cells (Tregs) (**Fig. 4c,d** and **fig. S3b**). Similarly, IL-12-encoding EGFR-LV effectively suppressed tumor growth in B16-EGFR5 and MC38-EGFR5 mouse models, demonstrating generalizability across different TAAs and tumor types (**Fig. 4e,f** and **fig. S3c,d**).

**Fig. 4.**
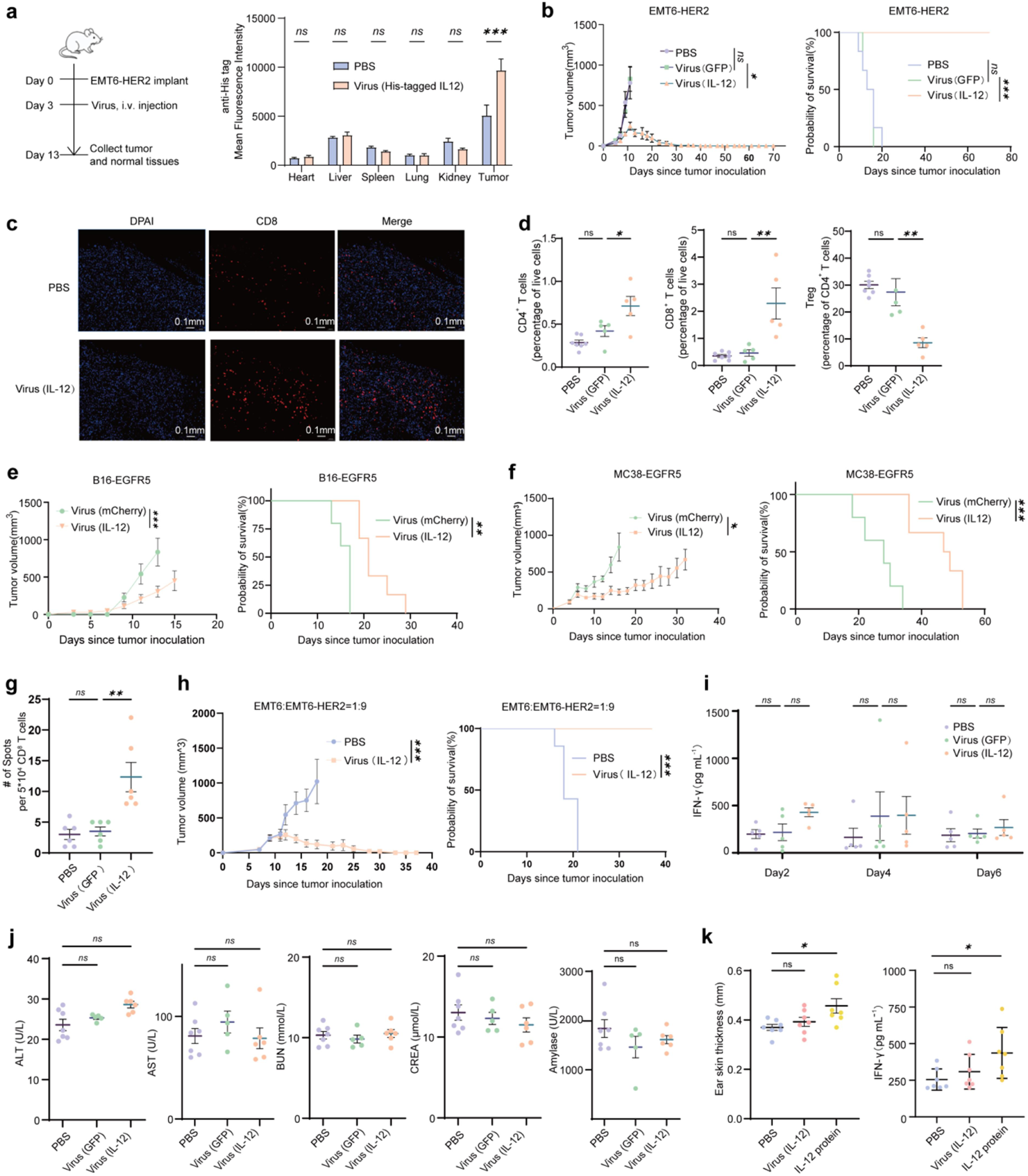
Tumor-restricted IL-12 production via LENTRA. (a) Mice bearing EMT6-HER2 tumors received PBS or His-tagged IL-12-encoding HER2-LV. After ten days, single-cell suspensions prepared from tumors and normal tissues were stained with Alexa Fluor™ 647 anti-His tag antibody, and analyzed by flow cytometry. (b) Tumor growth and survival of EMT6-HER2-bearing mice treated with PBS (n = 6), GFP-encoding HER2-LV (n = 6), or IL-12-encoding HER2-LV (n = 10). (c) Mice bearing EMT6-HER2 tumors were treated with PBS, or IL-12-encoding HER2-LV. After nine days, tumors were sectioned, stained with anti-CD8a antibody, and imaged (scale bar, 0.1 mm). (d) Mice bearing EMT6-HER2 tumors were treated with PBS, GFP-encoding HER2-LV, or IL-12-encoding HER2-LV. After nine days, tumors were dissociated and analyzed by flow cytometry for immune cell populations. (e) Tumor growth and survival of mice bearing B16-EGFR5 tumors treated with mCherry- or IL-12-encoding EGFR-LV (n=5). (f) Tumor growth and survival of mice bearing MC38-EGFR5 tumors treated with mCherry- or IL-12-encoding EGFR-LV (n=5). (g) Mice bearing EMT6-HER2 tumors were treated with PBS, GFP-encoding HER2-LV, or IL-12-encoding HER2-LV. After nine days, CD8⁺ T cells isolated from spleens were co-cultured with parental EMT6 cells and analyzed by IFNγ ELISpot. (h) Tumor growth and survival of mice bearing heterogeneous tumors (90% EMT6-HER2 + 10% EMT6) treated with PBS (n=6) or IL-12-encoding HER2-LV (n=10). (i) Mice bearing EMT6-HER2 tumors were treated with PBS, GFP-encoding HER2-LV, or IL-12-encoding HER2-LV. Serum IFNγ levels were measured 2, 4, and 6 days post-treatment. (j) Serum clinical chemistry markers (ALT, AST, BUN, CREA, amylase) measured 6 days post-treatment. (k) DNFB-induced contact hypersensitivity mice bearing EMT6-HER2 tumors were treated intravenously with recombinant IL-12 protein or IL-12-encoding HER2-LV. Ear thickness and serum IFNγ levels were measured after DNFB challenge. ***p < 0.0001; **p < 0.01; *p < 0.05 by Student’s t-test for (a, g); by log-rank (Mantel-Cox) test for survival curves in (b, e, f, h); by one-way ANOVA with Tukey’s post-hoc test for (d, i, j, k); and by two-way ANOVA with Tukey’s post-hoc test for tumor growth curves in (b, e, f, h).

The reconfigured TME potently primed systemic antitumor immunity. ELISpot analysis showed that splenocytes from Lenti-IL-12-treated mice—but not Lenti-GFP controls—mounted a robust, antigen-specific IFNγ response upon *ex vivo* restimulation with EMT6 cells (**Fig. 4g**), indicating successful activation and expansion of endogenous tumor-reactive T cell clones. Beyond T cell priming, this locally initiated immunity also exerted a bystander effect that overcame antigenic heterogeneity. In tumors composed of 90% HER2-positive and 10% HER2-negative cells, HER2-targeted Lenti-IL-12 remained highly effective, achieving complete response rates comparable to those in homogeneous HER2⁺ tumors (**Fig. 4h** and **fig. S3e**). Thus, the endogenous immune response ignited by IL-12-encoding HER2-LV can eliminate antigen-loss variants, addressing a key limitation of therapies that depend on direct antigen engagement.

Despite this efficacy, tumor-restricted IL-12 production using HER2-LV did not cause severe systemic toxicity. Serum IFNγ levels (**Fig. 4i**) and markers of liver and kidney function (ALT, AST, BUN, CREA, and amylase) were comparable between Lenti-IL-12–treated mice and untreated controls (**Fig. 4j**). In the DNFB-induced contact hypersensitivity model(27), Lenti-IL-12–treated mice exhibited ear swelling and serum IFNγ levels similar to the PBS group, whereas systemic administration of recombinant IL-12 protein significantly exacerbated both parameters (**Fig. 4k**). These results confirm the safety advantage of HER2-LV in confining cytokine activity to the TME.

### *In situ* sialidase production via LENTRA remodels the TME

Having shown the antitumor efficacy of tumor-localized cytokine production, we asked whether LENTRA could be applied to therapeutic enzymes that remodel cell surface components, rather than simply binding cell surface receptors. We engineered tumor-targeted lentiviral vectors encoding sialidase, an enzyme that removes terminal sialic acid residues from glycoproteins and glycolipids on tumor and stromal cells. Pioneering studies from Bertozzi, Wu, and Läubli have shown that targeted tumor desialylation using antibody- or bispecific antibody-fused sialidases can induce tumor control *in vivo*(32–35). This activity involves two complementary mechanisms: removal of hypersialylated glycans that engage inhibitory Siglec receptors on myeloid and lymphoid cells(32, 36), and reduction of the steric and electrostatic barrier imposed by the tumor glycocalyx(34, 37). Rather than delivering a pre-made enzyme, LENTRA turns tumor cells into local factories that produce sialidase on site. We reasoned that tumor-restricted sialidase expression would establish a sustained, localized desialylation program within the TME.

A single intravenous dose of HER2-targeted lentivirus encoding His-tagged *Vibrio cholerae* sialidase (Lenti-Sia) yielded detectable enzyme expression in orthotopic EMT6-HER2 tumors, with no expression in normal tissues and no overt systemic toxicity (**Fig. 5a,b** and **fig. S4a**), confirming tumor-restricted payload expression.

**Fig. 5.**
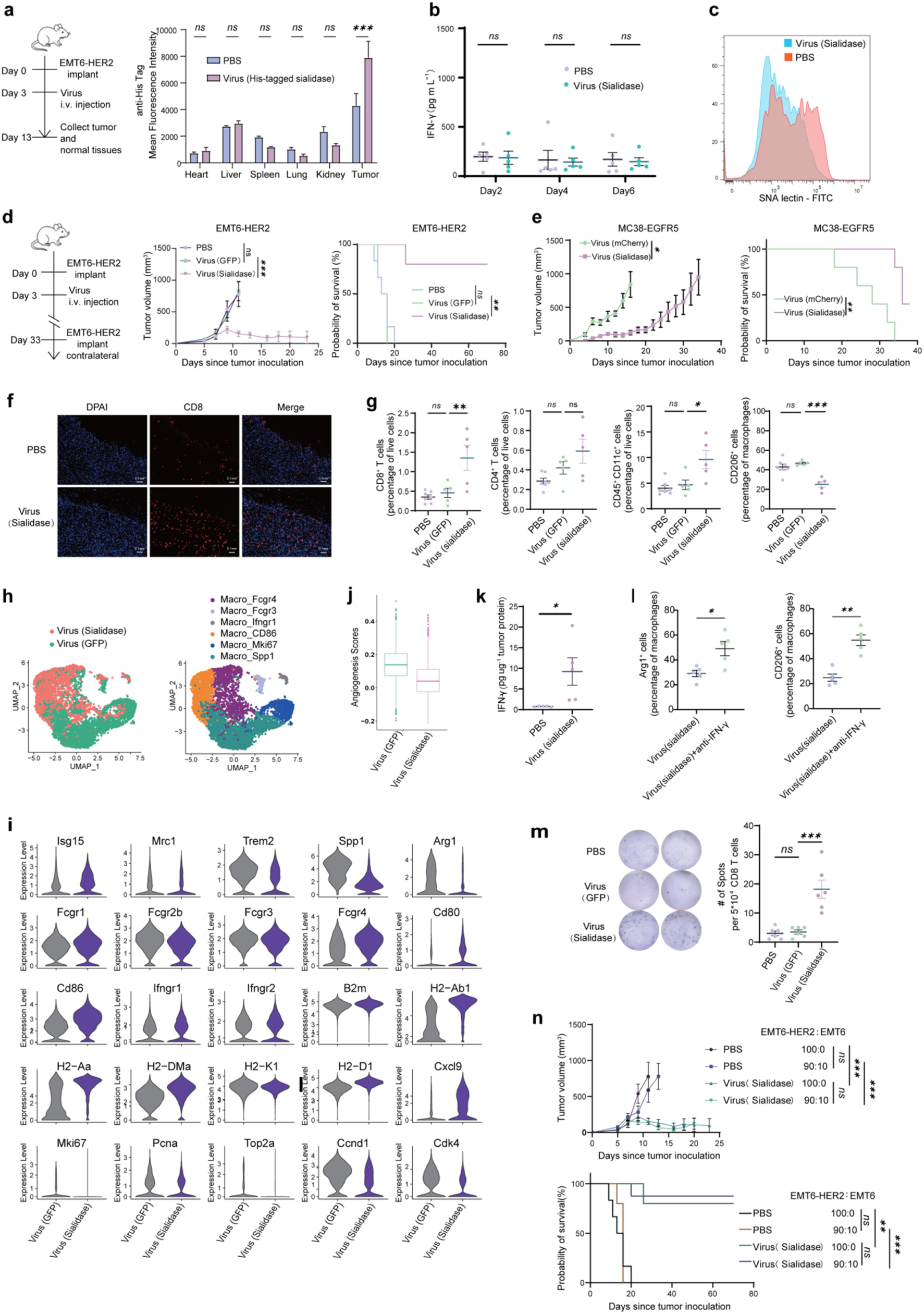
*In situ* sialidase production via LENTRA remodels the TME. (a) Mice bearing EMT6-HER2 tumors were treated with PBS or His-tagged sialidase-encoding HER2-targeting lentivirus (Lenti-Sia). Ten days later, single-cell suspensions prepared from tumors and normal tissues were stained with Alexa Fluor™ 647-conjugated anti-His tag antibody and analyzed by flow cytometry. (b) Mice bearing EMT6-HER2 tumors were treated with PBS or Lenti-Sia. Serum IFN-γ levels were measured by ELISA. (c) Tumors from (a) were dissociated, stained with SNA lectin-FITC, and analyzed by flow cytometry. (d) Tumor growth and survival of mice bearing EMT6-HER2 tumors treated with PBS (n=6), GFP-encoding HER2-LV (n=6), or Lenti-Sia (n=5). (e) Tumor growth and survival of mice bearing MC38-EGFR tumors treated with mCherry- or sialidase-encoding EGFR-LV (n=5). (f) Mice bearing EMT6-HER2 tumors were treated with PBS or Lenti-Sia. Tumors were sectioned, stained for CD8α, and imaged (scale bar, 100 μm). (g) Mice bearing EMT6-HER2 tumors were treated with PBS, GFP-encoding HER2-LV, or Lenti-Sia. Tumors were dissociated and analyzed by flow cytometry for immune cell populations. (h) UMAP projections showing TAM clusters from Lenti-GFP- or Lenti-Sia-treated tumors, annotated based on signature gene expression. (i) Violin plots showing differential expression of signature genes in TAMs from Lenti-GFP- versus Lenti-Sia-treated tumors. (j) Angiogenesis scores of TAMs from Lenti-GFP- or Lenti-Sia-treated tumors. (k) Mice bearing EMT6-HER2 tumors were treated with PBS or Lenti-Sia. Tumors were homogenized and the IFN-γ levels were analyzed by ELISA. (l) Mice bearing EMT6-HER2 tumors were treated with Lenti-Sia alone or in combination with an IFN-γ blocking antibody. Ten days later, tumors were harvested, dissociated, and analyzed by flow cytometry for CD206 and Arg1 expression on TAMs. (m) Mice bearing EMT6-HER2 tumors were treated with PBS, GFP-encoding HER2-LV, or Lenti-Sia. After nine days, CD8⁺ T cells isolated from spleen were co-cultured with EMT6 cells and analyzed by IFNγ ELISpot. (n) Tumor growth and survival of mice bearing homogeneous EMT6-HER2 tumors or heterogeneous tumors (90% EMT6-HER2 + 10% EMT6) treated with PBS (n=5-6) or Lenti-Sia (n=5-8). ***p < 0.0001; **p < 0.01; *p < 0.05 by Student’s t-test for (a, b, k, l); by log-rank (Mantel-Cox) test for survival curves in (d, e, n); by one-way ANOVA with Tukey’s post-hoc test for (g, m); and by two-way ANOVA with Tukey’s post-hoc test for tumor growth curve in (d, e, n).

*Vibrio cholerae* sialidase is a broad-spectrum enzyme that hydrolyzes terminal α2,3-, α2,6-, and α2,8-linked sialic acids from glycoproteins and glycolipids(32, 38). Lenti-Sia treatment markedly reduced tumor-cell surface sialylation (**Fig. 5c**), suppressed tumor growth, induced complete responses in 80% of treated mice, and extended survival in the EMT6-HER2 model (**Fig. 5d** and **fig. S4b**). Mice that achieved complete remission rejected a contralateral rechallenge with EMT6-HER2 tumor cells, indicating durable systemic immune memory (**Fig. 5d** and fig. S4b). Similarly, EGFR-targeted Lenti-Sia showed therapeutic activity in the subcutaneous MC38-EGFR model (**Fig. 5e** and **fig. S4c**).

To assess the impact of local sialidase on the TME, mice bearing EMT6-HER2 tumors were treated with HER2-targeted Lenti-Sia or PBS. Immunohistochemistry and flow cytometry of tumors showed that Lenti-Sia remodeled the immune landscape, with increased CD8⁺ T and dendritic cell infiltration, and reduced CD206⁺ M2-like macrophages (**Fig. 5f–g** and **fig. S4d**). To gain a higher-resolution view, we performed single-cell RNA sequencing of tumor-infiltrating immune cells. Unsupervised clustering showed substantial transcriptional reprogramming of tumor-associated macrophages (TAMs), with TAMs from Lenti-Sia-treated tumors forming clusters that were separate from control TAMs (**Fig. 5h**). Lenti-Sia treatment was associated with near-complete depletion of pro-tumorigenic clusters (Macro_Spp1 and Macro_Mki67) and emergence of immunostimulatory clusters (Macro_CD86 and Macro_Fcgr4) (**Fig. 5h**). Given that SPP1⁺ macrophages correlate with poor prognosis across multiple human cancers(39), their selective reduction suggests that localized desialylation using Lenti-Sia shifts the myeloid compartment toward a therapeutically favorable state.

Consistent with this, TAMs from Lenti-Sia-treated tumors showed elevated expression of T-cell chemoattractants (Cxcl9), antigen-presentation machinery (H2-Ab1, H2-Aa, H2-K1, H2-D1), and co-stimulatory molecules (Cd80, Cd86) (**Fig. 5i**). These TAMs also showed reduced expression of pro-tumorigenic and immunosuppressive signatures (Spp1, Arg1, Fcgr2b) and lower angiogenic activity compared with controls (**Fig. 5i,j**).

Lenti-Sia thus converts immunologically “cold” tumors into inflamed lesions, consistent with the increased intratumoral IFN-γ levels observed after treatment (**Fig. 5k**). To test whether IFN-γ signaling helps sustain the immunostimulatory TAM state, we treated EMT6-HER2 tumor-bearing mice with Lenti-Sia alone or in combination with IFN-γ blockade. Blocking IFN-γ reversed Lenti-Sia-induced myeloid remodeling, increasing CD206 and Arg1 expression in TAMs (**Fig. 5l**), suggesting that IFN-γ signaling contributes to maintaining the immunostimulatory phenotype after desialylation.

Lenti-Sia-mediated TME remodeling also promoted systemic antitumor immunity. ELISpot analysis showed that splenocytes from Lenti-Sia-treated mice, but not Lenti-GFP controls, mounted a tumor-specific IFN-γ response upon *ex vivo* restimulation with parental EMT6 cells (**Fig. 5m**), indicating successful priming of endogenous tumor-reactive T cells. Moreover, Lenti-Sia remained effective against antigenically heterogeneous tumors. In tumors composed of 90% EMT6-HER2 and 10% HER2-negative EMT6 cells, Lenti-Sia induced complete responses in 80% of mice, comparable to efficacy in homogeneous HER2-positive tumors (**Fig. 5n** and **fig. S4e**). Thus, sialidase-driven TME remodeling promotes bystander tumor clearance and antigen spreading, enabling elimination of antigen-negative variants that would otherwise escape targeted therapies.

### Translational validation in patient-derived tumor explants

After validating tumor-restricted transduction in mice, we asked whether LENTRA could also transduce and reprogram human tumors. We tested this using patient-derived tumor fragments, which preserve the architecture and cellular diversity of the original TME(40) (**Fig. 6a**). EGFR-positive tumor biopsies were dissected into fragments of approximately 1 mm^3^ and incubated with mCherry-encoding EGFR-LV. Unlike primary human immune cells, which were resistant to infection (**fig. S5a**), tumor cells were efficiently transduced, as shown by mCherry expression in dissociated fragments 5 days post-infection (**Fig. 6b**).

**Fig. 6.**
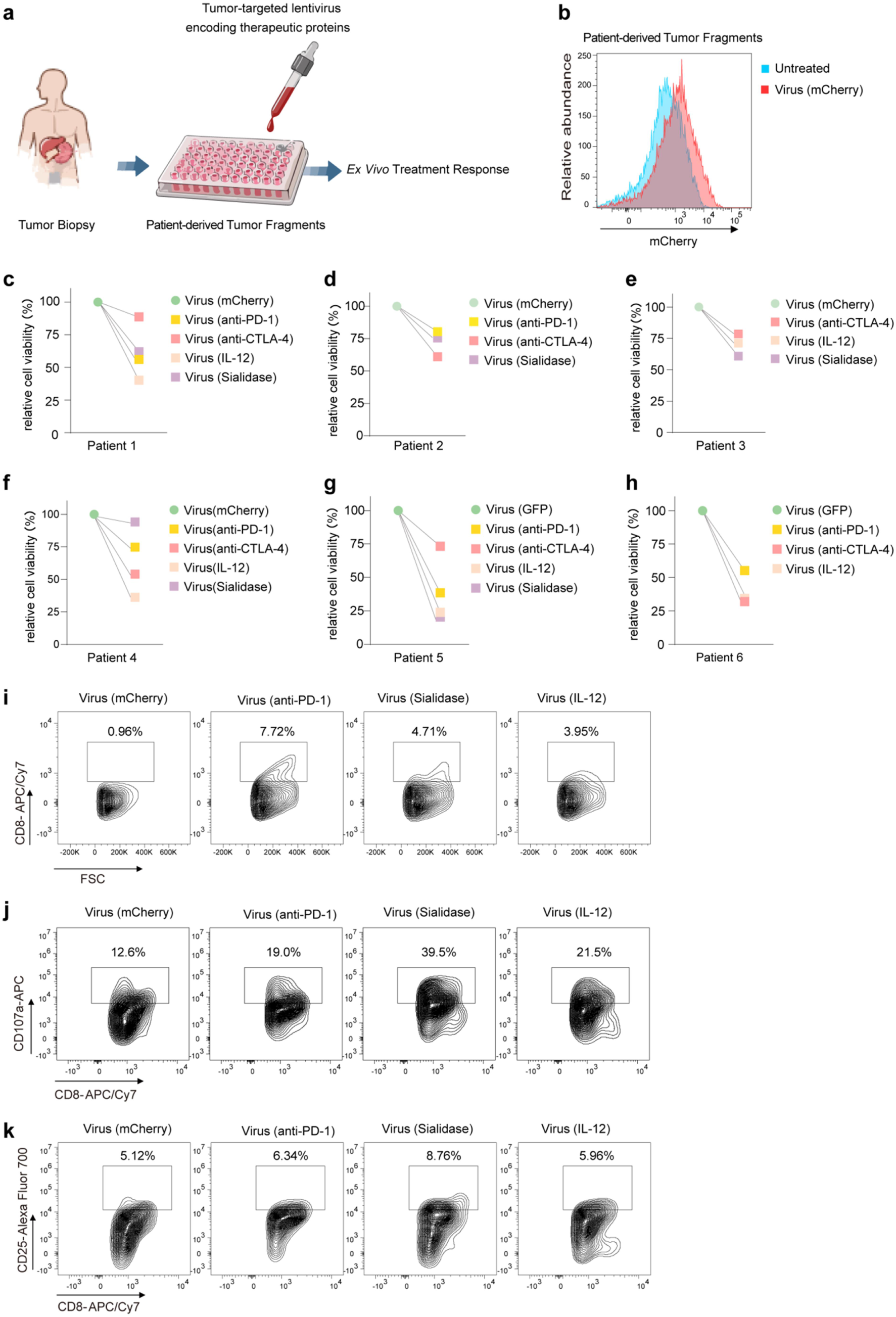
Translational validation in patient-derived tumor explants. (a) Experimental workflow for patient-derived tumor explants. (b) EGFR⁺ patient-derived tumor explants were treated with mCherry-encoding EGFR-LV. Single-cell suspensions were analyzed by flow cytometry for mCherry expression on tumor cells after 5 days. (c-h) Patient-derived EGFR⁺ (c-f) or HER2⁺ (g,h) tumor fragments were treated with the indicated lentivirus for 5 days. Cell viability was measured by CCK-8 assay and normalized to Lenti-mCherry or Lenti-GFP controls (set at 100%). (i-k) Patient-derived EGFR^+^ tumor fragments were treated with the indicated lentivirus for 5 days. Tumor-infiltrating CD8⁺ T cells were analyzed by flow cytometry for frequency (i), CD107a expression (j), and CD25 expression (k).

We then assessed antitumor activity. EGFR⁺ tumor fragments were treated *ex vivo* with EGFR-targeted lentivirus encoding anti-PD-1, anti-CTLA-4, IL-12, or sialidase. Viability assays 5 days later showed significant reduction in viable tumor cells across all payload groups relative to GFP controls (**Fig. 6c-f**). Comparable effects were seen in HER2⁺ tumor fragments treated with the corresponding HER2-targeted vectors (**Fig. 6g,h**).

Beyond direct killing, these vectors also remodeled the TME. In EGFR⁺ liver cancer fragments from patient 1, EGFR-LV increased intratumoral CD8⁺ T cell frequency from 0.96% (Lenti-mCherry) to 7.72% (Lenti-anti-PD-1), 3.95% (Lenti-IL-12), and 4.71% (Lenti-sialidase) (**Fig. 6i**). Anti-PD-1- and IL-12-encoding EGFR-LV upregulated CD107a on tumor-infiltrating CD8⁺ T cells, suggesting enhanced cytotoxic degranulation (**Fig. 6j**). Sialidase-encoding EGFR-LV increased both CD25 and CD107a on these cells (**Fig. 6j,k**). A similar pattern emerged in HER2⁺ breast cancer fragments from patient 5: compared with Lenti-GFP controls, Lenti-anti-PD-1 or Lenti-IL-12 increased CD107a and/or CD25 expression (**fig. S5b,c)**, while Lenti-sialidase increased CD8⁺ T cell frequency (**fig. S5d**), indicating distinct but convergent modes of T cell activation. Thus, LENTRA can transduce primary human tumors, reprogram them into *in situ* biofactories, and elicit both direct antitumor effects and local immune remodeling—supporting the translational potential of this approach for human solid tumors.

## Discussion

An ideal cancer therapy would act locally at the tumor site without exposing the rest of the body to toxic agents. Here we show that LENTRA, a tumor antigen-targeted lentiviral system, can specifically reprogram solid tumors into localized sources of therapeutic proteins, achieving potent efficacy in patient-derived tumor explants and robust tumor regression in multiple murine models after systemic delivery. By confining production of checkpoint inhibitors, cytokines, or enzymes to the TME, LENTRA allows the use of agents otherwise limited by systemic toxicity.

This approach also redefines the therapeutic role of tumor cells. Rather than treating them solely as immune targets, LENTRA repurposes tumor cells as local producers of immunomodulatory payloads. The tumor itself becomes the source of the immunostimulatory signal that orchestrates its own destruction, establishing a self-targeting circuit: tumor-derived payloads remodel the TME, recruit endogenous immunity, and drive clearance.

LENTRA offers a complementary strategy to emerging *in vivo* CAR-T therapies. Both bypass *ex vivo* manufacturing hurdles, but they approach solid tumors differently. *In vivo* CAR-T relies on engineering circulating T cells that must overcome exhaustion, infiltration barriers, and systemic activation risks(20, 41). By contrast, engineering tumor cells creates a localized, tumor-dependent source of therapy that persists as long as transduced cells remain. This may be especially valuable in immunologically “cold” tumors, where success requires not only effector activation but also broad TME remodeling.

Achieving high specificity remains a central challenge for systemic viral delivery. Many adeno-associated virus (AAV) vectors show broad tissue tropism and hepatic accumulation after intravenous administration(42). They also carry a packaging limit of ∼4.7 kb, restricting payload size and complexity(43). Lentiviral vectors support larger payloads and integrate into the host genome, enabling stable expression in proliferating tumor cells. LENTRA further enhances specificity through a decoupled targeting design, which separates tumor antigen recognition from membrane fusion via a VSV-G mutant and minimizes off-target transduction.

Our approach has several limitations. First, effective targeting depends on sufficient tumor-associated antigen expression, and heterogeneity may affect transduction efficiency. However, as our heterogeneous tumor models show, local immune activation can elicit bystander effects that eliminate antigen-negative cells. Second, despite improved specificity, off-target transduction and long-term safety require thorough evaluation before clinical translation. Future designs could incorporate regulatable expression systems to control payload production. Finally, permanent genetic modification carries a low but non-zero risk of insertional mutagenesis. This could be addressed by including a safety switch, such as inducible Caspase-9(44), which allows rapid apoptosis of transduced cells upon pharmacological trigger. Alternatively, LENTRA could be adapted to an integrase-deficient lentiviral vector (IDLV) format, delivering payloads as non-integrating episomes and eliminating insertional oncogenesis risk. While IDLV-mediated expression is transient due to episomal dilution in dividing cells, this could be advantageous for exceptionally potent payloads, such as IL-12, where a short, acute burst may suffice to prime antitumor immunity.

The LENTRA platform is modular. Both the targeting moiety and therapeutic payload can be exchanged, enabling adaptation to different tumor antigens and treatment strategies. For example, future applications could adapt LENTRA to bispecific engagers or prodrug-converting enzymes. Additionally, lentiviral vectors are already produced under established GMP-compliant pipelines, which may facilitate translation. Thus, LENTRA represents a clinically tractable strategy for programming tumors to produce therapeutic proteins locally within the TME.

## Materials and Methods

### Cell Lines

HEK293T, MDA-MB-231, MDA-MB-453, SK-BR-3, and EMT6 cells were obtained from the Institute of Biochemistry and Cell Biology, Chinese Academy of Sciences (Shanghai, China). HEK293T, MDA-MB-231, and MDA-MB-453 cells were cultured in Dulbecco’s Modified Eagle’s Medium (DMEM; Gibco) supplemented with 10% fetal bovine serum (FBS; Sigma) and 1% penicillin-streptomycin (Thermo Fisher). SK-BR-3 and EMT6 cells were maintained in RPMI 1640 medium (Gibco) supplemented with 10% FBS and 1% penicillin-streptomycin. All cell lines were incubated at 37°C in a humidified atmosphere containing 5% CO₂.

MC38-HER2, MC38-EGFR5, EMT6-HER2, EMT6-EGFR5, B16-HER2, and B16-EGFR5 cell lines were generated by lentiviral transduction. The plasmid pLVX-HER2 was constructed by subcloning HER2-mCherry (HER2 cDNA fused to mCherry) cDNA into the lentiviral vector pLVX. Lentiviruses were produced by transient transfection of HEK293T cells using 40 kDa linear polyethylenimine (PEI; Yeasen) at a PEI:DNA mass ratio of 1.5:1. The plasmid mass ratio for transfer plasmid, packaging plasmid (psPAX2), and fusogen plasmid (VSV-G) was 3:2:1, respectively. Lentiviruses were concentrated and used to transduce MC38, EMT6, and B16 cells. Transduced cells were sorted by FACS two days post-infection to isolate populations with high mCherry expression. Similarly, EMT6-EGFR5, MC38-EGFR5, and B16-EGFR5 cells were generated by transducing parental lines with lentivirus encoding an EGFR-mEGFP fusion construct, followed by FACS sorting for high mEGFP expression. The plasmid pLVX-EGFR5 was constructed by subcloning cDNA of a chimeric mouse EGFR with six amino acid mutations followed by mCherry cDNA into the lentiviral vector pLVX.

### Plasmid cloning and construction

Primers were synthesized by GENEWIZ. The plasmid pMD2.G (Addgene #12259) was digested with EcoRI to remove the wild-type VSV-G fragment, which was replaced with a mutated VSV-G fragment (K47Q, R354A) generated by PCR primers to create the VSV-G double mutant. Targeting plasmids for lentivirus production were constructed based on the pMD2 backbone. The HER2- or EGFR-targeting plasmid was generated by inserting a gene fragment encoding an anti-HER2 nanobody (clone: 2Rs15d) or anti-EGFR scFv (derived from cetuximab), together with a CD8a stalk linker and CD8a transmembrane domain, into pMD2. cDNAs encoding Vibrio cholerae sialidase (UniProt: P0C6E9), single-chain IL-12 (p40 and p35 linked by a (G₄S)₃ linker), anti-CTLA-4 scFv (clone: 9D9), and anti-PD-1 scFv (clone: RMP1-14) were synthesized by GENEWIZ. Transfer plasmids were generated by inserting the IL-2 signal sequence together with the cDNA of sialidase, single-chain IL-12, anti-CTLA-4 scFv, or anti-PD-1 scFv, followed by a 6XHis tag, into a lentiviral vector.

### Lentivirus production

Lentiviruses were produced by transient transfection of HEK293T cells using 40 kDa linear PEI (Yeasen) at a PEI:DNA mass ratio of 1.5:1. Briefly, DNA and PEI were diluted separately in DMEM, mixed, and incubated at room temperature for 30 min. The complexes were added dropwise to cells. After 4-6 h, the medium was replaced with DMEM containing 10% FBS and 1% penicillin-streptomycin. The plasmid mass ratio for transfer plasmid, packaging plasmid (psPAX2), targeting plasmid, and fusogen plasmid (mutant or wild-type VSV-G) was 3:2:1.5:1, respectively. Lentiviral supernatants were collected 48 h post-transfection and centrifuged at 4,500 rpm for 10 min at 4°C. Supernatants were either used directly or concentrated using Lenti-X Concentrator (Takara) according to the manufacturer’s protocol. For concentration, virus was precipitated at 4,500 rpm for 60 min at 4°C, and the pellet was resuspended in PBS and stored at -80°C. For infection, 2 μL of concentrated lentivirus was added to 50,000 target cells in a 24-well plate. Transduction efficiency was assessed by flow cytometry 48 h post-infection.

### Testing the selectivity of HER2-targeted lentivirus in mixed cell populations

MDA-MB-231 cells were labeled with CellTrace dyes (CFSE or Cell Proliferation Dye eFluor 450; Thermo Fisher) according to the manufacturer’s instructions and mixed with MDA-MB-453 cells at specified ratios (1:1, 1:10, 1:100, or 1:1000). Concentrated HER2-targeted lentivirus (3 μL) was added to 50,000 cells in a 24-well plate; alternatively, 12 μL was added to 200,000 cells in a 6-well plate. After 48 h, cells were washed and analyzed by flow cytometry to determine infection efficiency in CellTrace-positive and -negative populations, or imaged using a confocal laser scanning microscope (Nikon, Japan).

### HER2-targeted lentivirus infectivity in cells with variable HER2 surface density

To generate cells with varying HER2 surface density, HEK293T cells were transduced with increasing volumes (2, 6, 10, 12, or 16 μL per 50,000 cells in a 24-well plate) of HER2-mCherry-encoding lentivirus. After 48 h, these cells were transduced with 2 μL of concentrated GFP-encoding HER2-targeted lentivirus. After an additional 48 h, cells were analyzed by flow cytometry to determine transduction efficiency.

### HER2-targeted lentivirus infectivity in the presence of HER2-blocking antibody

MDA-MB-453 cells were incubated with anti-HER2 antibody at indicated concentrations (0, 0.78, 3.91, 19.55, 97.75, or 163 nM) for 10 min at room temperature. After two washes with PBS, 4 μL of concentrated HER2-targeted lentivirus in 400 μL DMEM was added. The medium was replaced with fresh complete DMEM after 8 h. Cells were analyzed by flow cytometry 48 h post-infection.

### *In vivo* selectivity of HER2- or EGFR-targeted lentivirus

C57BL/6 and BALB/c mice (6 weeks old) were purchased from Gempharmatech Co., Ltd. (Nanjing, China). All animal procedures were approved by the institutional animal care and use committee and conducted in accordance with institutional guidelines.

EMT6 or EMT6-HER2 tumors were established by orthotopic implantation of 3 × 10⁵ EMT6-HER2 cells into the mammary fat pad of six-week-old female BALB/c mice. Three days after tumor implantation, mice received an intravenous injection of 3 × 10⁶ TU of GFP-encoding HER2-LV (in 100 μL PBS) or PBS. Ten days later, tumors and major organs (heart, liver, spleen, lung, kidney) were harvested. GFP signals were detected using an IVIS

Spectrum CT system (PerkinElmer) with excitation and emission filters set at 500 nm and 540 nm, respectively. Tissues were mechanically dissociated and passed through a 70-μm cell strainer using a syringe plunger to obtain single-cell suspensions. Cells were stained with Fixable Viability Dye eFluor™ 450 (Thermo Fisher) to exclude dead cells and analyzed by flow cytometry for GFP expression.

For the B16-EGFR5 tumor model, six-week-old C57BL/6 mice were subcutaneously inoculated with 3 × 10⁵ B16-EGFR5 cells. Six days later, mice received an intravenous injection of 3 × 10⁶ TU of mCherry-encoding EGFR-targeted lentivirus (in 100 μL PBS). Nine days after viral administration, tumors were harvested and processed as described above; mCherry expression was analyzed by flow cytometry.

To evaluate anti-CTLA-4, anti-PD-1, sialidase, or IL-12 expression, EMT6-HER2 tumor-bearing mice (established as above) were treated intravenously with 3 × 10⁶ TU (in 100 μL PBS) of HER2-LV encoding 6XHis-tagged anti-CTLA-4, anti-PD-1, sialidase, or IL-12, or with PBS three days after tumor implantation. Ten days later, tumors and major organs were harvested and processed into single-cell suspensions. Cells were stained with Fixable Viability Dye eFluor™ 450, washed, and Fc receptors were blocked with purified anti-mouse CD16/32 antibody (BioLegend) for 10 min at room temperature. Cells were then surface-stained with anti-CD45 antibody (clone 30-F11, BioLegend) for 30 min at 4°C. Intracellular staining was performed using the Cytofix/Cytoperm™ Fixation/Permeabilization Kit (BD Biosciences) according to the manufacturer’s instructions, followed by staining with anti-His-tag antibody (clone J095G46, BioLegend) for 30 min at 4°C. Cells were analyzed by flow cytometry.

### Quantitative PCR analysis

Genomic DNA was extracted from organs and tumor tissues using the Universal Genomic DNA Purification Mini Spin Kit (Beyotime) according to the manufacturer’s instructions. Total RNA was extracted using TRIzol reagent (Thermo Fisher); RNA concentration and purity were determined using a NanoDrop spectrophotometer (Thermo Fisher). First-strand cDNA was synthesized from 1 μg total RNA using a BeyoRT™ III First Strand cDNA Synthesis Kit (Beyotime) with random hexamer primers. Quantitative PCR was performed using Hieff® qPCR SYBR® Green Master Mix (Yeasen) on a Real-Time PCR System (Bio-Rad). Reaction conditions were: 95°C for 5 min, followed by 40 cycles of 95°C for 10 s and 60°C for 30 s. Gene expression levels were normalized to *Gapdh*, and relative quantification was calculated using the 2⁻ΔΔCt method. Primer sequences were:

*mEGFP:*

Forward: 5′–GAACCGCATCGAGCTGAA–3′

Reverse: 5′–TGCTTGTCGGCCATGATATAG–3′

*Gapdh:*

Forward: 5′–ACGGCCGCATCTTCTTGTGCA–3′

Reverse: 5′–ACGGCCAAATCCGTTCACACC–3′

Data are expressed as mean ± SD of three replicates. Statistical significance was assessed using two-tailed Student’s t-test.

### Mouse tumor models and treatment

EMT6-HER2 tumors were established by orthotopic implantation of 3 × 10⁵ EMT6-HER2 cells into the mammary fat pad of six-week-old female BALB/c mice. Three days later, mice received an intravenous injection of 3 × 10⁶ TU (in 100 μL PBS) of HER2-LV encoding GFP, IL-12, sialidase, anti-CTLA-4, or anti-PD-1. In parallel experiments, EMT6-EGFR5 tumor-bearing mice received EGFR-LV encoding mCherry, anti-CTLA-4, or anti-PD-1 on the same schedule.

For the B16-EGFR5 model, six-week-old C57BL/6 mice were subcutaneously inoculated with 3 × 10⁵ B16-EGFR5 cells and received an intravenous injection of 3 × 10⁶ TU (in 100 μL PBS) of EGFR-LV encoding mCherry, IL-12, anti-CTLA-4, or anti-PD-1 six days later.

For the MC38 model, C57BL/6 mice were subcutaneously inoculated with 5 × 10⁵ MC38-EGFR5 cells and received an intravenous injection of 3 × 10⁶ TU (in 100 μL PBS) of EGFR-LV encoding mCherry, IL-12, or sialidase four days later.

### Analysis of tumor-infiltrating immune cells

EMT6-HER2 tumors were established by orthotopic implantation of 3 × 10⁵ EMT6-HER2 cells into the mammary fat pad of six-week-old female BALB/c mice. Three days later, mice were treated with PBS (control) or 3 × 10⁶ TU (in 100 μL PBS) of IL-12- or sialidase-encoding HER2-LV. On day 12, tumors were harvested, weighed, and mechanically dissociated. Single-cell suspensions were prepared by filtering through 70-μm cell strainers. Cells were stained with Fixable Viability Dye eFluor™ 450, washed, and Fc receptors were blocked with anti-mouse CD16/32 antibody (BioLegend) for 10 min at room temperature. Cells were then stained with the following antibodies at 4°C for 30 min: CD4-FITC (clone RM4-5), CD8α-APC (clone 53-6.7), CD11c-FITC (clone N418), I-A/I-E-APC/Cy7 (clone M5/114.15.2), CD80-PE (clone 16-10A1), CD11b-APC/Cy7 (clone M1/70), F4/80-FITC (clone BM8), CD206-PE (clone C068C2), CD45-APC (clone 30-F11), and CD25-PE (clone PC61) (all from BioLegend). Stained cells were analyzed by flow cytometry.

### Immunofluorescence imaging of mouse tissue sections

EMT6-HER2 tumors were established by orthotopic implantation of 3 × 10⁵ EMT6-HER2 cells into the mammary fat pad of six-week-old female BALB/c mice. Three days later, mice were treated with PBS (control) or 3 × 10⁶ TU (in 100 μL PBS) of sialidase-encoding HER2-LV. On day 12, tumors were harvested, fixed in 4% paraformaldehyde overnight, and paraffin-embedded. Sections underwent antigen retrieval for 30 min, blocking with 3% BSA for 30 min, and incubation with anti-CD8a antibody (Servicebio) at 4°C overnight. After three washes, sections were incubated with secondary antibody (Servicebio) for 50 min at room temperature, washed, and stained with DAPI for 10 min. Slides were imaged using a digital slide scanner (3DHISTECH).

### Intratumoral IFN-γ analysis

EMT6-HER2 tumor-bearing mice were treated as described in the tumor-infiltrating immune cell analysis. Single-cell suspensions were prepared and adjusted to 10⁷ cells/mL in RPMI 1640 medium. A 100 μL aliquot was subjected to three freeze-thaw cycles (dry ice and 37°C water bath) to lyse cells. The lysate was centrifuged at 8,000 × g for 5 min, and the supernatant was collected. IFN-γ concentration was measured using the ELISA MAX™ Standard Set Mouse IFN-γ kit (BioLegend) according to the manufacturer’s instructions.

### ELISpot assay

EMT6-HER2 tumors were established by orthotopic implantation of 3 × 10⁵ EMT6-HER2 cells into the mammary fat pad of six-week-old female BALB/c mice. Three days later, mice were treated with PBS (control) or 3 × 10⁶ TU (in 100 μL PBS) of GFP- or sialidase-encoding HER2-LV. On day 12, spleens were harvested, mechanically dissociated, and filtered through 70-μm cell strainers. CD8⁺ T cells were isolated using the EasySep™ Mouse CD8⁺ T Cell Isolation Kit (STEMCELL). EMT6 cells were pretreated with 100 IU/mL murine IFN-γ (Sino Biological) for 20 h and washed three times with PBS. CD8⁺ T cells (5 × 10⁴) and EMT6 cells (1 × 10⁴) were co-cultured overnight on PVDF membranes pre-coated with anti-IFN-γ capture antibody (BioLegend) in 96-well plates. Plates were washed three times with PBS containing 0.1% Tween 20, incubated with biotinylated anti-IFN-γ detection antibody (BioLegend) for 1.5 h, washed, and incubated with streptavidin-alkaline phosphatase (Yeasen) for 1 h. After three washes, BCIP/NBT substrate was added. After 25 min, plates were rinsed with water, air-dried, and analyzed using an ELISpot reader (CTL).

### DNFB-induced contact hypersensitivity model

The dorsal skin of six-week-old female BALB/c mice was shaved and sensitized by topical application of 25 μL 1.0% DNFB (1-fluoro-2,4-dinitrobenzene; Aladdin) dissolved in a 4:1 mixture of acetone and olive oil. Five days after sensitization, mice were subcutaneously inoculated with 3 × 10⁵ EMT6-HER2 cells. Three days after tumor implantation, mice received 3 × 10⁶ TU (in 100 μL PBS) of IL-12- or anti-CTLA-4-encoding HER2-LV intravenously, 200 μg recombinant anti-CTLA-4 protein (BioXCell) intraperitoneally, or recombinant IL-12 protein intraperitoneally. Four and ten days later, mice were challenged by topical application of 0.2% DNFB to the ear. Ear thickness was measured 24 h after each challenge.

### Serum cytokine and biochemistry analysis

For serum IFN-γ analysis, EMT6-HER2 tumor-bearing mice were treated with HER2-LV encoding GFP, IL-12, or sialidase. Blood samples were collected 2, 4, and 6 days after treatment and centrifuged at 3,000 × g for 15 min to obtain plasma. IFN-γ concentrations were measured using the ELISA MAX™ Standard Set Mouse IFN-γ kit (BioLegend) according to the manufacturer’s protocol. For serum biochemistry, plasma samples were diluted fourfold with sterile water. Levels of alanine aminotransferase (ALT), aspartate aminotransferase (AST), amylase, blood urea nitrogen (BUN), and creatinine (CREA) were measured using commercially available kits (Meikang Biotechnology Co., Ltd., Ningbo, China) according to the manufacturer’s instructions.

### *Ex vivo* evaluation of patient-derived tumor explants

Human tumor biopsies were dissected into fragments of approximately 1 mm³ and cultured in RPMI 1640 medium (Thermo Fisher Scientific) supplemented with 1 mM sodium pyruvate (Sigma-Aldrich), 1× MEM non-essential amino acids (Sigma-Aldrich), 2 mM L-glutamine (Thermo Fisher Scientific), 10% fetal bovine serum (FBS), 3,000 U/mL recombinant human IL-2, and 1% penicillin-streptomycin. The fragments were incubated in a 96-well plate with 1 × 10⁶ TU of EGFR-LV or HER2-LV encoding mCherry, GFP, IL-12, anti-PD-1, or anti-CTLA-4. After five days, fragment viability was assessed using a CCK-8 assay kit (according to the manufacturer’s instructions). To analyze the phenotype of tumor-infiltrating T cells, tumor fragments were digested in RPMI 1640 medium supplemented with DNase and 1 mg/mL collagenase type IV (Sigma-Aldrich) for 30 minutes at 37 °C under slow rotation. The resulting single-cell suspensions were washed with PBS, filtered through a 150-μm mesh, and resuspended in 50 μL of PBS. Cells were stained with Fixable Viability Dye eFluor™ 450 (Thermo Fisher Scientific) to exclude dead cells, followed by Fc receptor blocking (BioLegend). Subsequently, cells were stained with APC-anti-human CD107a (BioLegend #328620), Alexa Fluor 700-anti-human CD25 (BioLegend #356118), and APC/Cy7-anti-human CD8a (BioLegend #300926) for 30 minutes at 4 °C. After two washes, cells were analyzed by flow cytometry.

### Single-cell RNA sequencing

EMT6-HER2 tumors were established by orthotopic inoculation of 3 × 10⁵ cells into female BALB/c mice. On day 3, mice were treated with 3 × 10⁶ TU (in 100 μL PBS) of sialidase- or GFP-encoding HER2-LV. Tumors were harvested on day 16 (13 days post-treatment). Tumors from nine mice per group were excised and minced into 2-3 mm fragments. Tumor tissues from all mice per group were pooled and thoroughly mixed to ensure uniform representation. Pooled tissues were enzymatically dissociated into single-cell suspensions, and CD45⁺ and CD45⁻ cells were sorted and mixed at a 9:1 ratio. Single-cell suspensions were loaded onto a 10x Chromium controller using the Chromium Single-Cell 3′Kit (v3) according to the manufacturer’s instructions. cDNA amplification and library construction were performed following standard protocols. Libraries were sequenced by LC-Bio Technology (Hangzhou, China) on an Illumina NovaSeq 6000 system (paired-end, 150 bp) at a minimum depth of 20,000 reads per cell.

Raw sequencing data were converted to FASTQ format using bcl2fastq software (v5.0.1). Reads were aligned to the reference genome and quantified using Cell Ranger (v7.0.0) with default parameters. The resulting gene expression matrix was loaded into Seurat (v4.1.0) for downstream analysis. Low-quality cells were filtered based on the following criteria: number of genes expressed per cell >500 and mitochondrial gene percentage <25%. Data were normalized using the LogNormalize method, and principal component analysis (PCA) was performed on the top 2,000 variable genes. The top 20 principal components were used for clustering with the FindClusters function and for visualization using UMAP. Marker genes for each cluster were identified using the FindAllMarkers function with the following criteria: expression in >10% of cells in the cluster, adjusted P value ≤ 0.01, and log₂ fold change ≥ 0.26. Clusters containing macrophages were selected for re-analysis using the same processing and clustering steps. Differential gene expression analysis between specified populations was performed using the FindMarkers function.

## Statistical analysis

All figures are representative of at least three experiments unless otherwise noted. All graphs report mean ± standard deviation (SD) values of biological replicates. The statistical significance of two-group comparisons was calculated using Student’s t-test. Animal survival was analyzed using Log-rank (Mantel-Cox) test. Multi-group comparisons were carried out using one-way ANOVA with Tukey’s multiple comparisons test. Experiments with repeated measures over a time course, such as tumor growth, were analyzed using a RM (repeated measures) two-way based on a general linear model.

## Data Availability

The single-cell RNA-seq data have been deposited at GEO: GSE329559

## Acknowledgments

This work was supported by National Natural Science Foundation of China (22277109 to T.Z.) and Yangtze River Delta Science and Technology Innovation Community Joint Research Project (BK20254009 to W.L.). We thank the Shared Instrumentation Core Facility, Hangzhou Institute of Medicine Chinese Academy of Sciences.

## Author Contributions

SW, QX, MW, WL, and TZ designed the study. SW, QX, MW, JD, QG, HM, WL and TZ performed the experiments and analyzed the data. SW, QX, MW, WL, and TZ wrote the manuscript.

