## Supplemental Figures for "Systemic tumor-cell reprogramming converts solid tumors into *in situ* biofactories for immunotherapy"

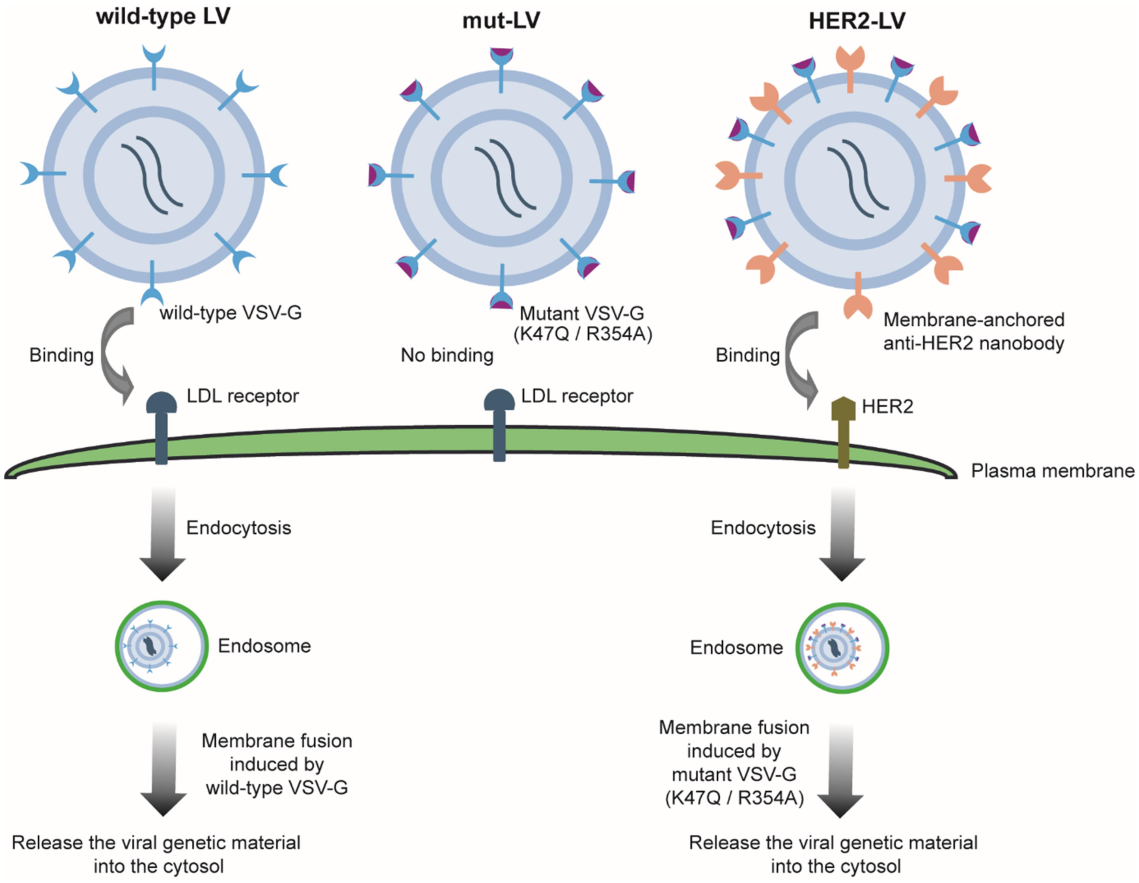


**Supplemental Figure S1.** Schematic illustration of the three lentiviral vectors used in this study. WT-LV is pseudotyped with wild-type VSV-G; mut-LV is pseudotyped with mutant VSV-G (K47Q/R354A) alone; HER2-LV is pseudotyped with both mutant VSV-G (K47Q/R354A) and a membrane-anchored anti-HER2 nanobody.

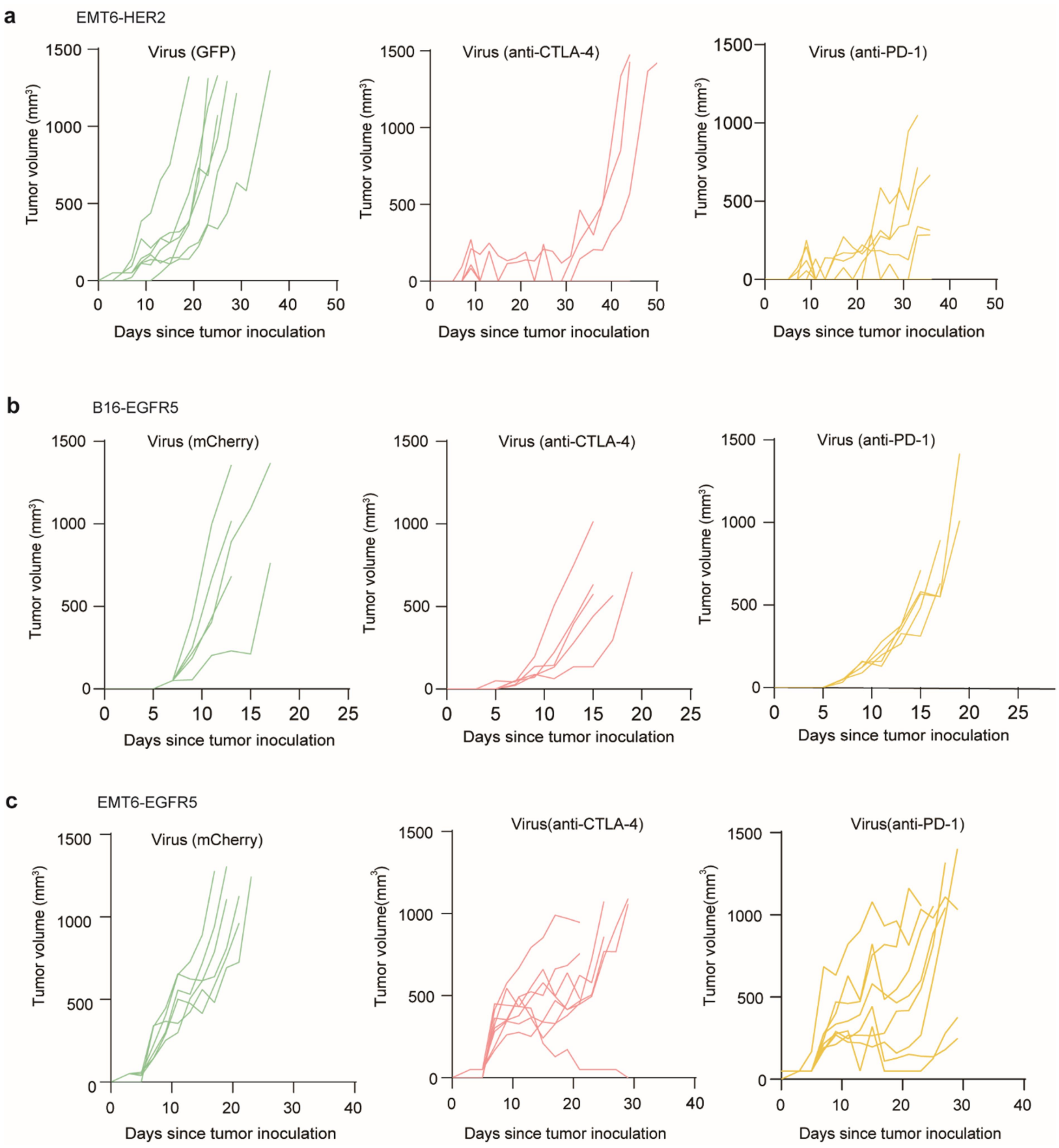


**Supplemental Figure S2.** LENTRA converts tumors into local factories for checkpoint blockade, related to Figure 3.

(a) Individual tumor growth curves of EMT6-HER2 tumor-bearing mice treated with HER2-targeted lentivirus encoding GFP, anti-CTLA-4, or anti-PD-1.

(b) Individual tumor growth curves of B16-EGFR5 tumor-bearing mice treated with EGFR-targeted lentivirus encoding mCherry, anti-CTLA-4, or anti-PD-1.

(c) Individual tumor growth curves of EMT6-EGFR5 tumor-bearing mice treated with EGFR-targeted lentivirus encoding mCherry, anti-CTLA-4, or anti-PD-1.

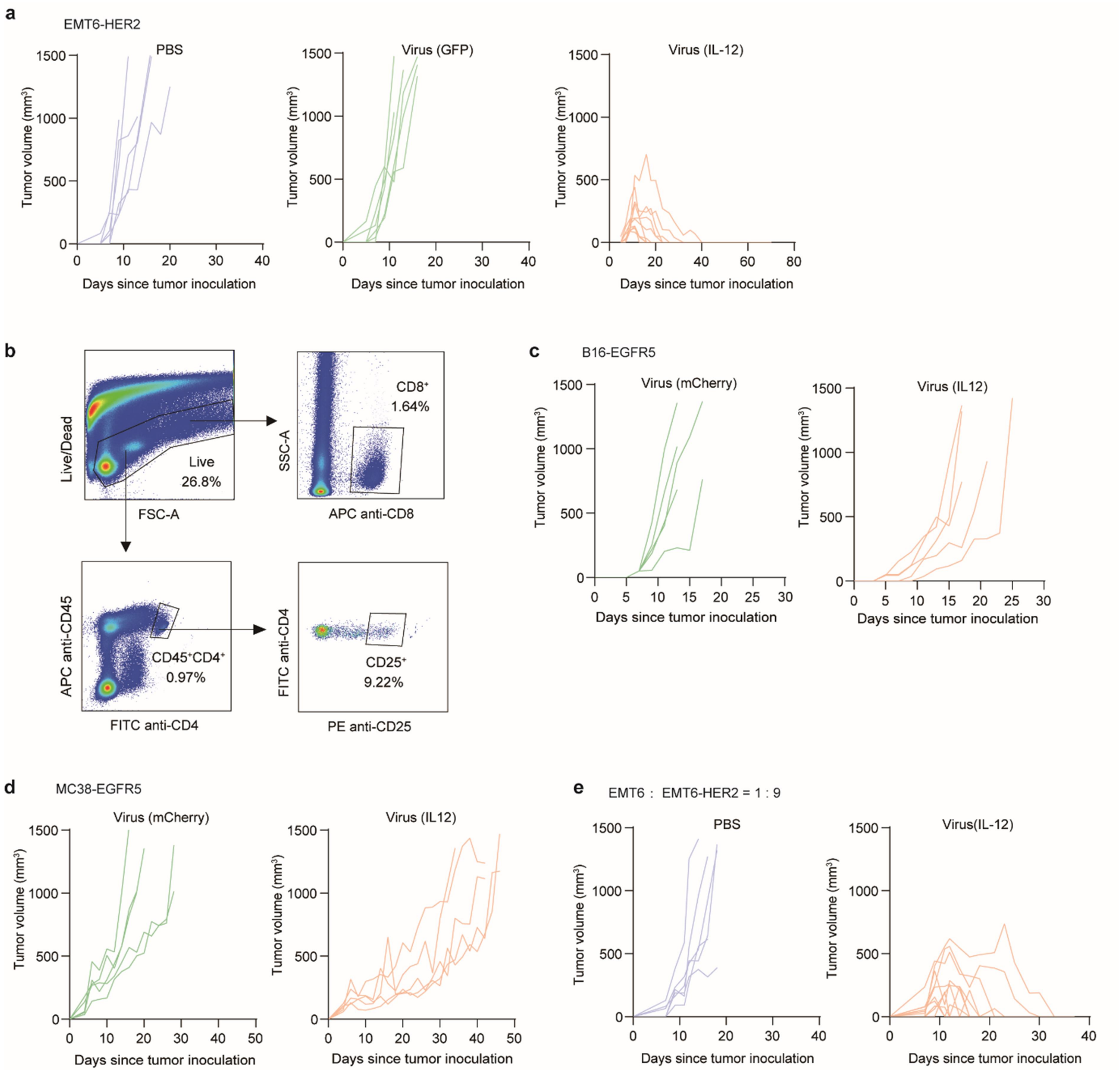


**Supplemental Figure S3.** Tumor-restricted IL-12 production via LENTRA, related to Figure 4.

(a) Individual tumor growth curves of EMT6-HER2 tumor-bearing mice treated with PBS, GFP-, or IL-12-encoding HER2-targeted lentivirus.

(b) Representative flow cytometry gating strategy for identification of $CD4^+$, $CD8^+$, and $CD4^+CD25^+$ T cells in EMT6-HER2 tumors.

(c) Individual tumor growth curves of B16-EGFR5 tumor-bearing mice treated with mCherry- or IL-12-encoding EGFR-targeted lentivirus.

(d) Individual tumor growth curves of MC38-EGFR5 tumor-bearing mice treated with mCherry- or IL-12-encoding EGFR-targeted lentivirus.

(e) Individual tumor growth curves of mice bearing heterogeneous tumors (EMT6 and EMT6-HER2 mixed at 1:9 ratio) treated with PBS, GFP-, or IL-12-encoding HER2-targeted lentivirus.

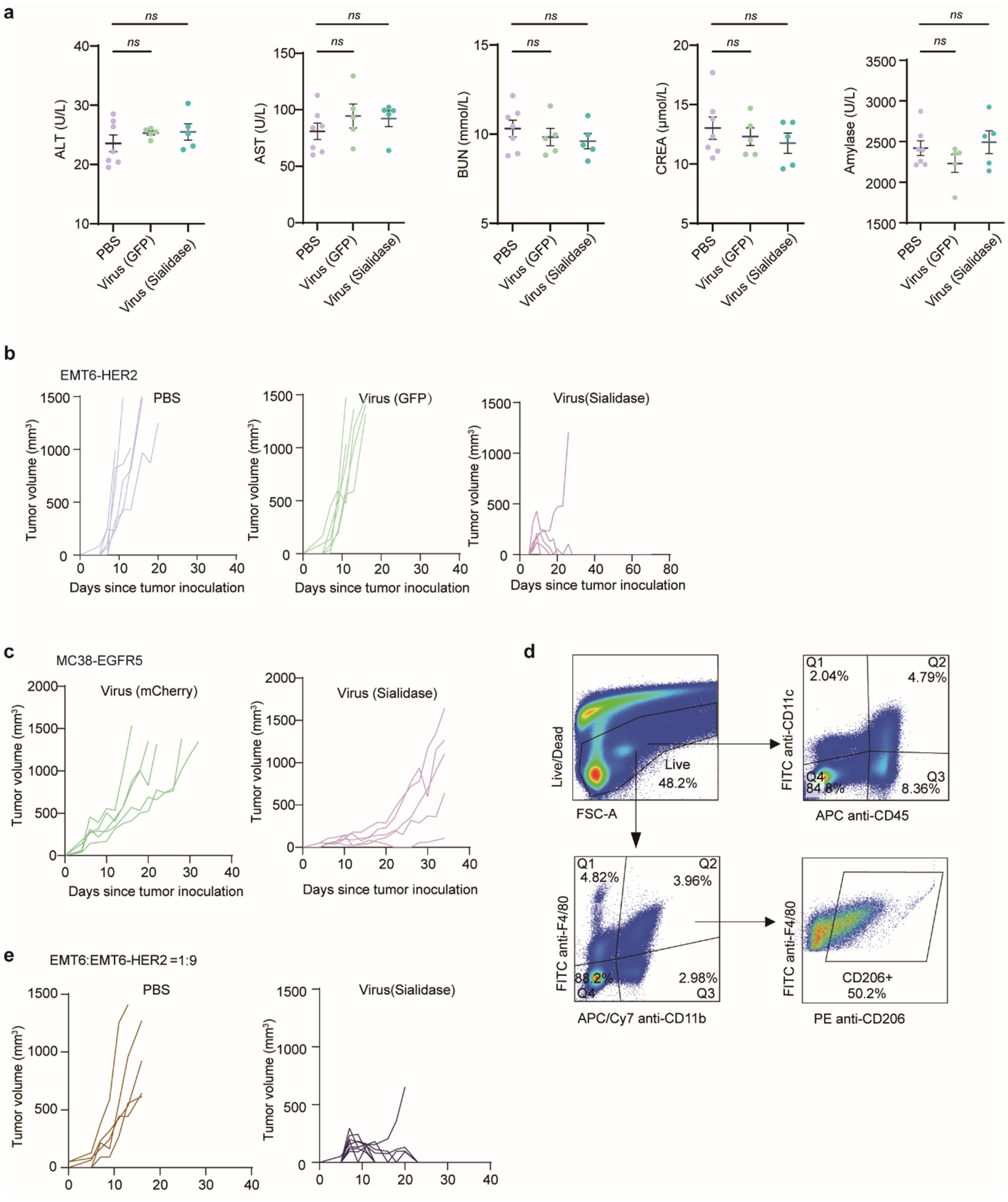


**Supplemental Figure S4.** *In situ* sialidase production via LENTRA remodels the TME, related to Figure 5.

(a) Serum levels of ALT, AST, BUN, CREA, and amylase in mice bearing EMT6-HER2 tumors treated with PBS, GFP- or sialidase-encoding HER2-LV.

(b) Individual tumor growth curves of EMT6-HER2 tumor-bearing mice treated with PBS, GFP- or sialidase-encoding HER2-targeted lentivirus.

(c) Individual tumor growth curves of MC38-EGFR5 tumor-bearing mice treated with mCherry- or sialidase-encoding EGFR-targeted lentivirus.

(d) Representative flow cytometry gating strategy for identification of $CD45^{+}CD11c^{+}$ dendritic cells and $F4/80^{+}CD11b^{+}CD206^{+}$ macrophages in EMT6-HER2 tumors.
(e) Individual tumor growth curves of mice bearing heterogeneous tumors (EMT6 and EMT6-HER2 mixed at 1:9 ratio) treated with PBS or sialidase-encoding HER2-targeted lentivirus
ns, not significant by one-way ANOVA with Tukey's post-test for (a).

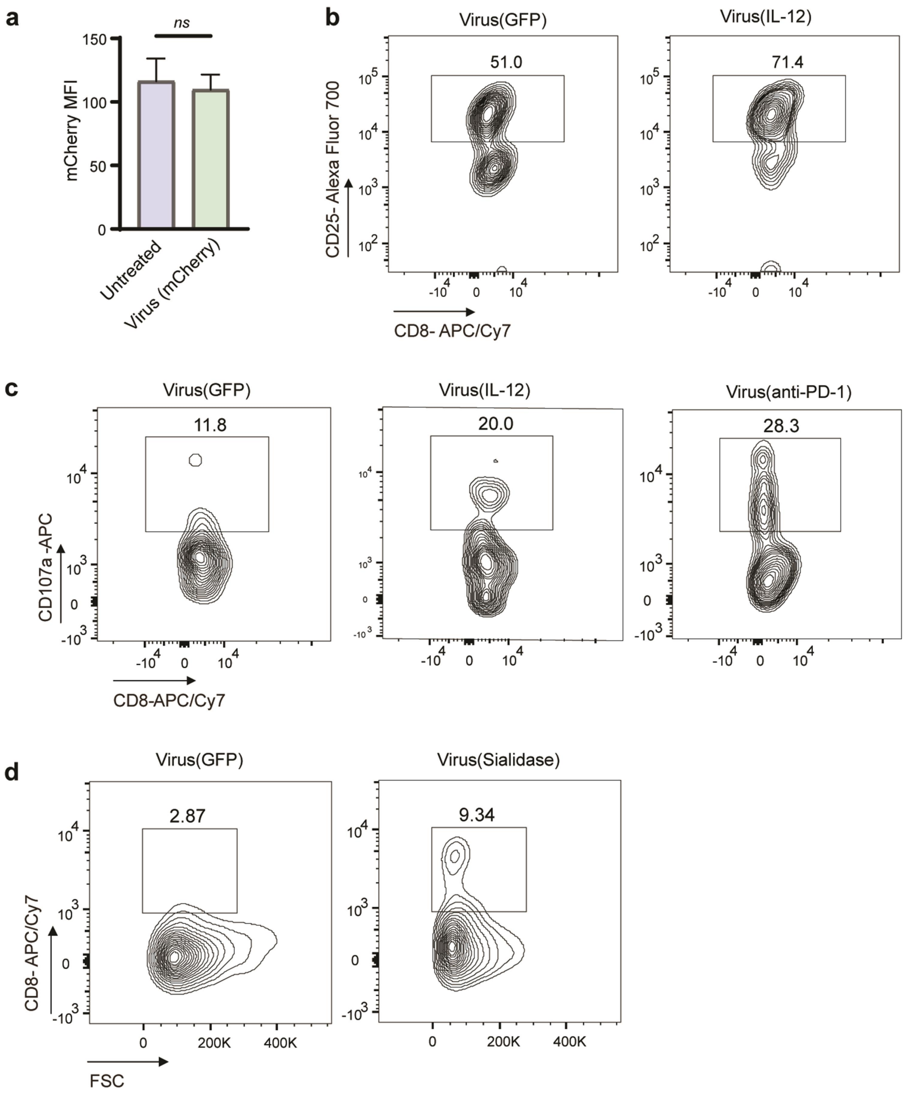


**Supplemental Figure S5.** Translational validation in patient-derived tumor explants, related to Figure 6.

(a) Human PBMCs were cultured in medium supplemented with mCherry-encoding EGFR-targeted lentivirus for 5 days, and mCherry expression was analyzed by flow cytometry.

(b–d) $HER2^+$ patient-derived tumor explants were treated with the indicated lentivirus for 5 days. Tumor-infiltrating $CD8^+$ T cells were analyzed by flow cytometry for (b) CD25 expression, (c) CD107a expression, and (d) frequency.

ns, not significant by Student's t-test for (a).
